# Neuraminidase activity modulates cellular co-infection during influenza A virus multicycle growth

**DOI:** 10.1101/2022.09.20.508375

**Authors:** Zijian Guo, Yuanyuan He, Ananya N. Benegal, Michael D. Vahey

## Abstract

Infection of individual cells by multiple virions plays critical roles in the replication and spread of many viruses, but mechanisms that control cellular co-infection during multi-cycle viral growth remain unclear. Here, we investigate virus-intrinsic factors that control cellular co-infection by influenza A virus (IAV). Using quantitative fluorescence to track the spread of virions from single infected cells, we identify the IAV surface protein neuraminidase (NA) as a key determinant of cellular co-infection. We map this effect to NA’s ability to deplete viral receptors from both infected and neighboring uninfected cells. In cases where viral infectious potential is low, genetic or pharmacological inhibition of NA increases the local spread of infection by increasing the viral load received by neighboring cells. These results identify virus-intrinsic factors that contribute to cellular multiplicity of infection, and suggest that optimal levels of NA activity depend on the infectious potential of the virus in question.

## Introduction

Single cells infected by influenza A virus (IAV) can give rise to hundreds to thousands of new virions within a single replication cycle^1–3^. These virions spread non-uniformly, producing wide variations in viral load that are concentrated around the initial site of infection. This phenomenon is widely recognized *in vitro*, providing the foundation for plaque assays for detecting and quantifying influenza and many other viruses. More recently, the development of reporter viruses^4–6^ and genetic barcoding strategies^7,8^ have revealed that viral spread *in vivo* exhibits similar features: for example, showing sensitivity to anatomical compartmentalization^9,10^.

An important consequence of non-uniform viral spread is that it produces wide variation in the cellular multiplicity of infection (MOI). This has implications that are particularly important in the biology of IAV. The IAV genome is comprised of eight distinct RNA segments, and although most virions fail to deliver all eight segments individually^11,12^, complementation through co-infection can sustain productive infection^3^. Cellular co-infection also enables reassortment when it occurs between distinct viral lineages^13^; it enhances the replication of viruses that are poorly adapted to their hosts^14^; and it can modulate cellular immune responses^15,16^. These wide-ranging contributions to influenza replication and spread make understanding the causes and consequences of IAV co-infection a high priority. However, the extent to which co-infection depends on specific properties of the virus or the target cell remains unclear.

Multiple factors may contribute to the spatial structure of viral spread and the degree of co-infection that occurs during multi-cycle growth. These include the number of virions released from infected cells (*i.e*., the burst size); physical association between virions as they transit the extracellular environment; and the typical distance traveled by virions before they attach to and enter a naïve cell (*i.e*., the degree of dispersal). Viral burst size can influence the frequency of co-infection by increasing the number of viral particles in the extracellular environment. Alternatively, viral aggregation^17^, bacterial hitchhiking^18,19^, and clustered packaging into extracellular vesicles^20^ have each been shown to enhance co-infection through the physical association of multiple infectious units. Finally, virion dispersal could influence the degree of co-infection by determining whether progeny virions remain concentrated at the site of infection or if they spread out over a large range. While each of these considerations could influence the spread of IAV, specific links between IAV genotype and co-infection through these different routes have not been identified.

In the case of viral dispersal, the IAV surface proteins hemagglutinin (HA) and neuraminidase (NA) are likely to contribute^21–23^. HA and NA are expressed on the surface of infected cells and packaged into virions, where they exhibit competing biochemical activities. NA cleaves the viral receptor, sialic acid, allowing virus particles to spread throughout the host^24,25^. Released virions can initiate a new round of cellular infection through HA-mediated attachment to sialic acid on the surface of a naïve cell^26^. While both HA and NA are essential for virus replication and transmission, evolutionary data and laboratory experiments demonstrate that consequential mutations to one protein can be tolerated through compensatory mutations to the other^27–31^. Biochemical data comparing HA avidity and NA catalytic activity for strains circulating in humans further supports the idea that efficient replication and transmission within a particular host depends on balance between the two proteins’ competing activities^32,33^. However, the precise consequences of imbalanced HA and NA on viral spread - and whether other factors contribute to how well imbalance is tolerated - is not well understood. Further complicating this picture, some NAs can bind to sialic acid^34^, and NAs cleave sialic acid both in *cis* (i.e., on the membrane of the infected cell)^35^ and in *trans* (as in a standard enzyme-linked lectin assay^36^), raising the question of how these activities collectively contribute to virus release and attachment.

To begin addressing these questions, we investigated mechanistic links between the degree of co-infection that occurs during multi-cycle IAV growth, and the *in situ* activity of HA and NA on cells and viruses. Using quantitative measurements of virion spread, we identify strain-specific differences in the degree of co-infection that different viruses support during multicycle growth in cell culture. We find that these differences are genetically linked to the NA segment, with the number of virions shed to neighboring cells varying inversely with NA enzymatic activity. We show that attenuation of NA activity using either genetic mutations or chemical inhibition can increase the spread of infection in viruses with high dependence on co-infection. Combined with stochastic modelling of virus spread, these results demonstrate the importance of viral adhesion and release in establishing the spatial structure of viral spread, and suggest that the optimal balance between HA and NA activities for a given strain will depend on its infectious potential

## Results

### An imaging-based approach to tracking virion spread

We developed a fluorescence-imaging based approach to monitor the local spread of single IAV virions, with the goal of determining the viral load per cell during secondary infection in units of physical particles. This method allows us to determine the degree of co-infection caused by viruses derived from a common lineage, which therefore lack defined genetic differences. Following infection at low MOI, we incubate cells with fluorescently-labeled Fab fragments that recognize conserved epitopes on HA away from the receptor binding site. This allows us to image, segment, and count individual virions shed from sites of infection without disrupting the ability of these virions to bind to sialic acid (Sia) receptors. By blocking viral fusion with the endosome, Fabs derived from CR9114^37^ and FI6v3^38^ restrict replication to a single round. From these measurements, we define local virion shedding as the number of virions originating from isolated sites of infection (typically one single infected cell) that bind to and/or are internalized by neighboring uninfected cells (Figure 1A). Volumetric imaging of infected A549 cells over time reveals that local virion shedding remains steady from 14-18 hours post-infection (Supplementary Figure 1); we therefore selected the ~16h timepoint for our data collection. Values for local virion shedding obtained in this way serve as a proxy for the degree of cellular co-infection that a particular viral strain supports during secondary infection.

**Figure 1:**
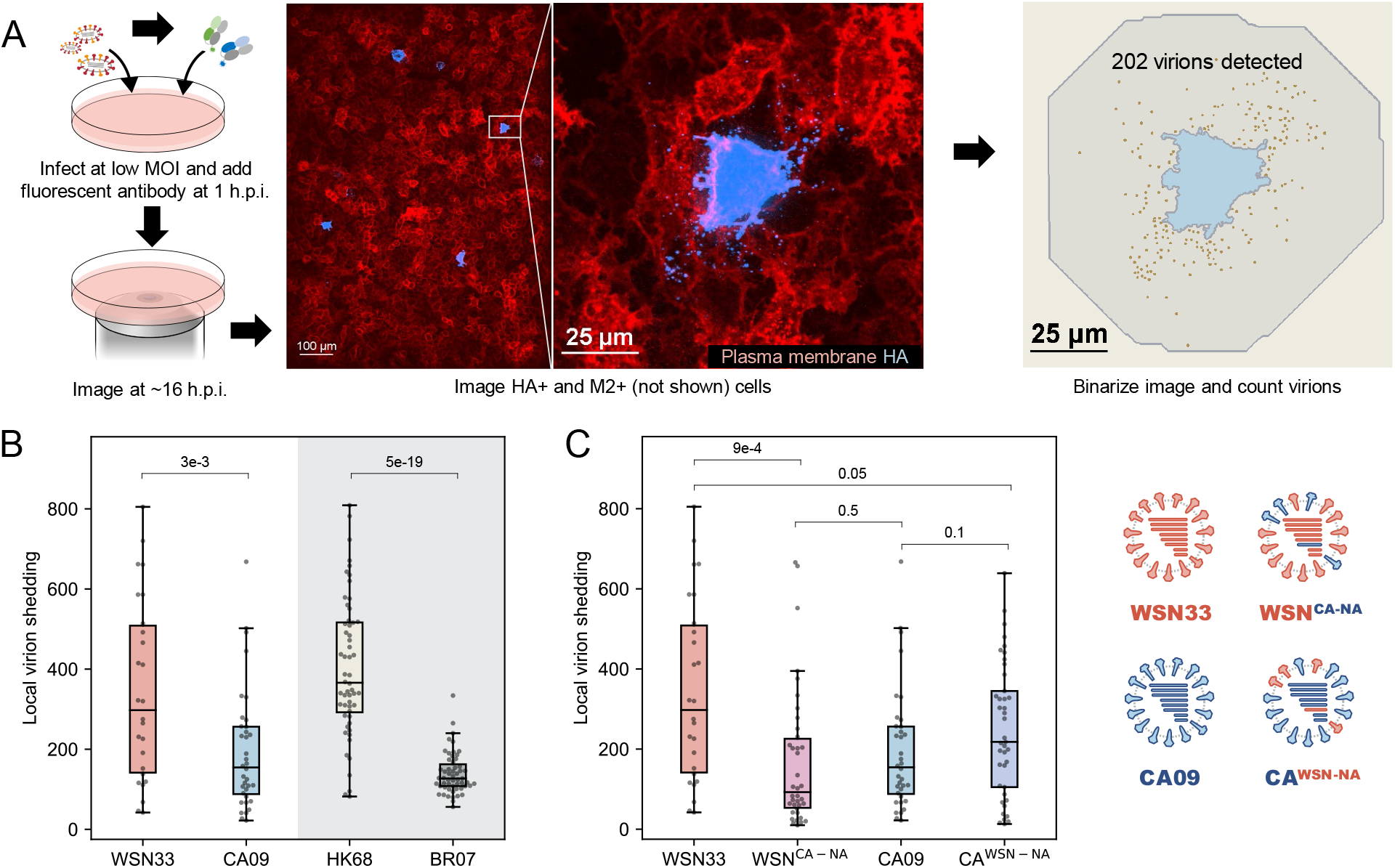
Virions produced by infected cells are preferentially shed to the nearest neighbors. (A) Schematic and representative images illustrating the approach for measuring local virion shedding. Contrast in the HA channel is exaggerated so that shed virions are visible. (B) Quantification of local virion shedding for four IAV strains. Data is combined from three biological replicates. Individual data points represent virions from a single site of infection. P-values are determined by independent t-tests. (C) Quantification of local virion shedding for WSN^CA-NA^ and CA^WSN-NA^. Wild-type results from (B) are shown for comparison. Data is combined from three biological replicates. P-values are determined by independent t-tests.

Using this approach, we first sought to measure the local spread of virions produced by different IAV strains and subtypes. We selected two H1N1 strains - A/WSN/1933 (WSN33) and A/California/04/2009 (CA09) - and two H3N2 strains - A/Hong Kong/1/1968 (HK68) and A/Brisbane/10/2007 (BR07). These viruses represent a combination of lab-adapted and contemporary strains for the two IAV subtypes currently circulating in humans. We find that the number of virions shed to neighboring uninfected cells differs markedly both within a single strain as well as between strains (Figure 1B), consistent with extreme heterogeneity in the outcome of IAV infection at low MOI^12^, and suggesting that genetic differences between strains and subtypes contribute to localized virion spread.

We hypothesized that the activity of neuraminidase (NA) may play an outsized role in establishing this phenotype. NA depletes cell-surface sialic acids^35^, reducing the chance of super-infection^39^ and promoting viral release^25^. To test if NA contributes to differences in local virion shedding observed between strains, we created recombinant viruses in which the NA segment of CA09 and WSN33 were exchanged. Expression of WSN33 NA in a CA09 genetic background increased local virion shedding (making it more WSN33-like), while expression of CA09 NA in a WSN33 background decreased local virion shedding (making it more CA09-like) (Figure 1C). These results establish local virion shedding as a viral phenotype that differs between strains and depends in part of the NA segment.

### *In situ* activity of NA differs between strains and shapes permissiveness to viral attachment

NA could contribute to local virion shedding in multiple ways: increasing or decreasing viral adhesion depending on its enzymatic activity, or influencing virion assembly through unrelated mechanisms. To help distinguish between these possibilities, we first sought to compare the ability of different NAs to deplete sialic acid from the cell surface. We labeled sialic acid on A549 monolayers infected at low MOI (~0.003) using mild periodate oxidation followed by conjugation with an aldehyde-reactive fluorophore^40^ (Supplementary Figure 2 & 3). Comparing Sia levels across cells infected with different strains provides a metric for the *in situ* activity of NA against its native substrates in cell culture. These measurements show that the selected strains deplete Sia to different extents (Figure 2A), and that the efficiency with which NA removes Sia from the cell surface does not generally correlate with its activity against MUNANA (Figure 2B), consistent with prior work evaluating NA activity against larger, multiply-sialylated substrates^41–43^.

**Figure 2:**
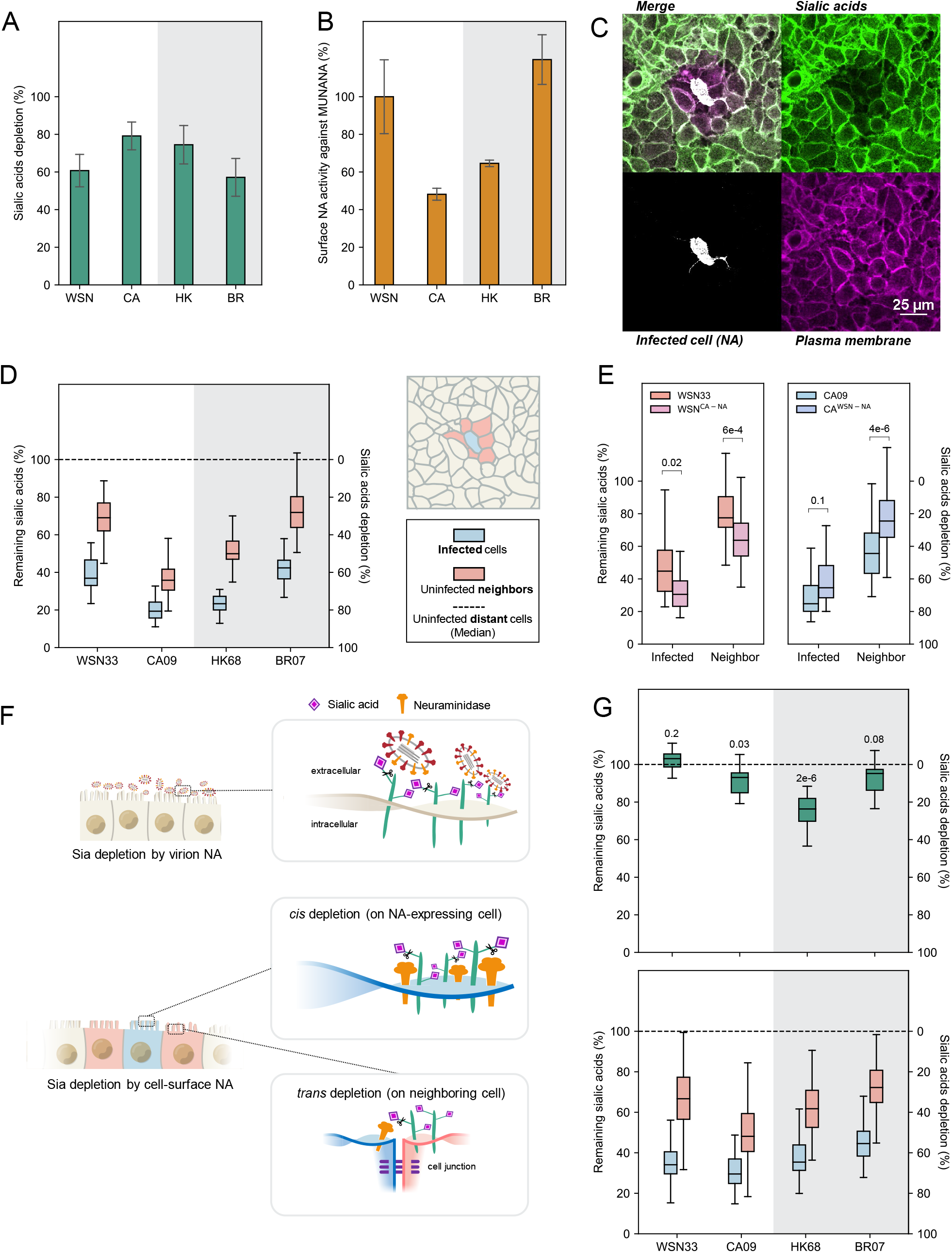
Cell-surface NA depletes sialic acid (Sia) in *cis* and in *trans*. (A) Cell-surface Sia depletion on cells infected by each viral strain. A value of 100% corresponds to complete depletion. (B) Surface NA activity against MUNANA, normalized to data for WSN33. (C) Image showing Sia, cell-surface NA, and plasma membrane in the proximity of an isolated cell infected with HK68. (D) Quantification of remaining Sia on the surface of infected cells and uninfected neighbors. Cells were selected based on NA expression by incubating with 1G01 immediately before imaging. Data is combined from three biological replicates. (E) Quantification of remaining Sia on the surface of infected cells (determined by HA expression) and uninfected neighbors for viruses with exchanged NA segments. Data is combined from three biological replicates. P-values are from independent t-tests. (F) Illustration of Sia depletion by virion-associated NA (top) and cell-associated NA (bottom). (G) *Top*: Quantification of Sia depletion following incubation with virions. Results are normalized to Sia levels on untreated cell surfaces. P-values are determined from independent t-tests relative to the untreated group. *Bottom*: Quantification of Sia depletion by cell-surface NA. Cells were selected based on NA expression. Data is combined from three biological replicates.

In addition to the depletion of Sia from the surface of infected cells (*i.e*., in *cis*), we also observe depletion from the surface of adjacent uninfected cells (*i.e*., in *trans*; Figure 2C). *Trans* depletion of Sia follows similar trends across the strains tested as *cis* depletion (Figure 2D), and also occurs in cells with greater apical-basal polarity (MDCK; Supplementary Figure 4). Viral strains with swapped NAs (WSN^CA-NA^ and CA^WSN-NA^) showed similar *cis* and *trans* Sia depletion as the strains from which their NAs were derived (Figure 2E). We reasoned that Sia depletion in *trans* could be driven by NA on the surface of cells, released viruses, or both. To evaluate the contributions of these two sources of NA activity, we compared Sia depletion driven by virus-associated NA with that of NA expressed on the surface of cells in the absence of other viral proteins (“cell-associated” NA) (Figure 2F). Viral NA (delivered to A549 monolayers at a density of ~10 virions per cell) shows a relatively modest effect on Sia depletion, reaching a maximum of ~25% for HK68 (Figure 2G, top), whereas cell-associated NA achieves similar *cis* and *trans* depletion of cell surface Sia as that observed in virus-infected cells (Figure 2G, bottom). Collectively, these results indicate that NA from virus-infected cells deplete Sia receptors both in *cis* and in *trans*, and that cell-surface NA in the absence of viral budding is sufficient to recapitulate trends observed during viral infection.

We next sought to determine how Sia depletion affects virion attachment for the four strains from our initial test. The HAs of these viruses have different avidity for human receptors^33,44^, suggesting that they may differ in their sensitivity to Sia depletion. To determine how attachment of each viral strain to A549 monolayers changes following reduction of cell surface Sia, we measured virus binding following treatment with exogenous sialidase (from *Clostridium perfringens; CpNA*) relative to an untreated control. As expected, the four strains respond differently to Sia depletion, with BR07 showing the greatest reduction even at modest levels of Sia depletion, and HK68 and CA09 showing the most persistent binding as Sia levels are reduced (Figure 3A). The level of Sia depletion achieved with exogenous *Cp*NA (~60%) is comparable to or less than that achieved through cell-surface expression of IAV NAs in the context of viral infection (~60-80% reduction; Figure 2A), suggesting that the *in situ* activity of NA would be sufficient to reduce virion attachment to background levels for these strains. Consistent with this prediction, we observe strong reductions in viral attachment in both NA-expressing cells as well as adjacent cells that do not express NA (Figure 3B). Taken together, these results demonstrate that viral HA influences the extent of virion attachment to cells, while *cellular* NA influences which cells are most permissive to attachment.

**Figure 3:**
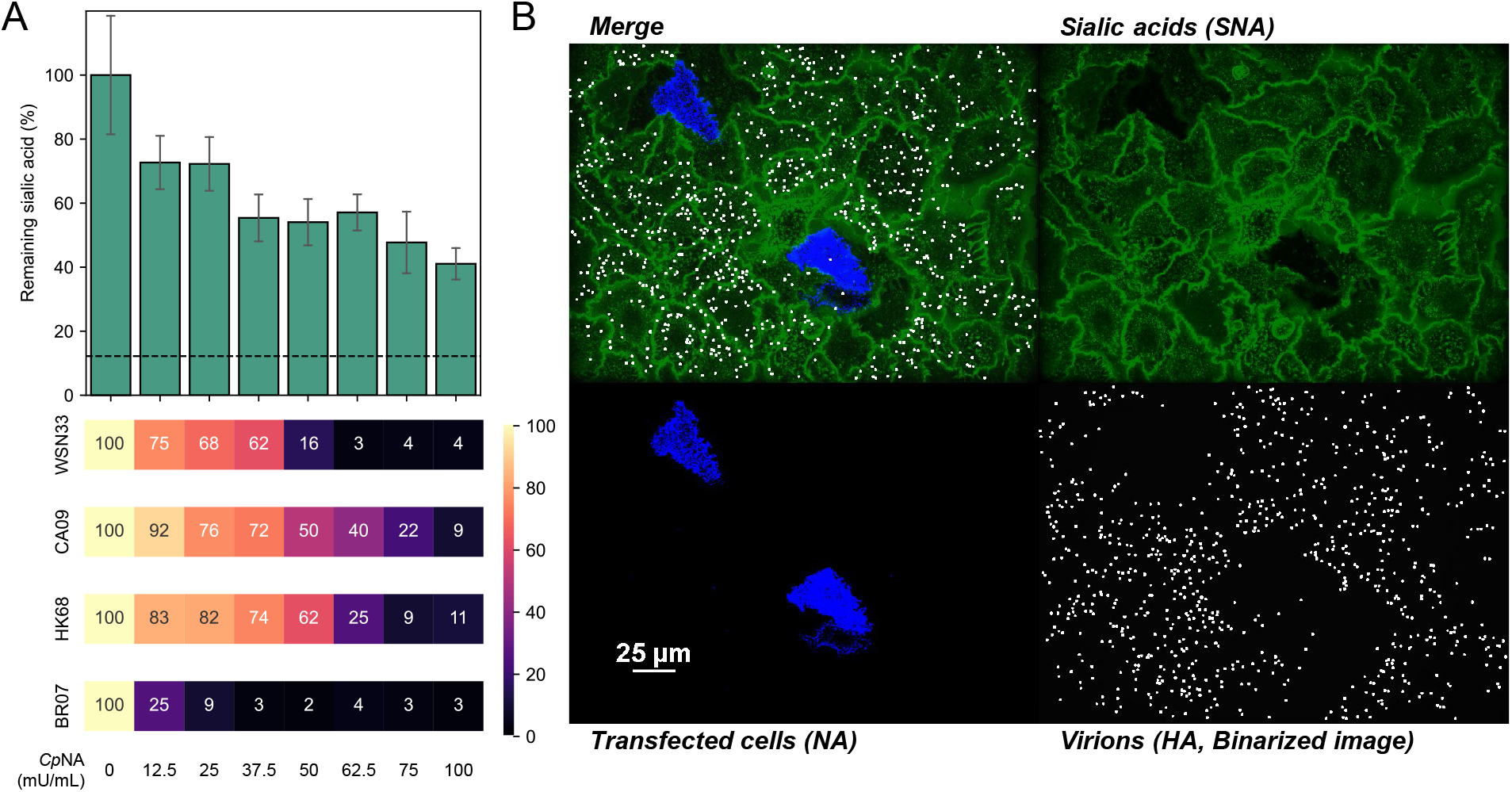
Sialic acid depletion reduces viral attachment in a strain-dependent manner. (A) *Top*: Cell surface Sia abundance following treatment with *Cp*NA; concentrations are specified below. Dashed line indicates signal from SLC35A1 knock-out cells treated with the highest *Cp*NA concentration. *Bottom*: Relative binding capacities for each viral strain following *Cp*NA treatment. Data is combined from three biological replicates. (B) Image showing virus attachment to cell monolayers with sparse expression of HK68 NA.

### Increased *in situ* NA activity reduces local cellular co-infection

Based on these observations, we reasoned that high *in situ* NA activity (as exhibited by CA09 NA; Figure 2A) could deplete Sia receptors from cells adjacent to the initial site of infection, reducing virus binding and uptake by these cells. This could potentially explain the differences in local virion shedding we observed when NA segments were swapped between WSN33 and CA09 (Figure 1C). To determine if direct cell-to-cell viral spread is sensitive to Sia depletion, we measured local virion shedding following treatment with exogenous sialidase (*Cp*NA). We observed a ~10-fold reduction in local virion shedding for WSN33-infected cells, and a ~5-fold reduction for cells infected with CA09 (Figure 4A), confirming that cell-to-cell viral spread is Sia-dependent.

**Figure 4:**
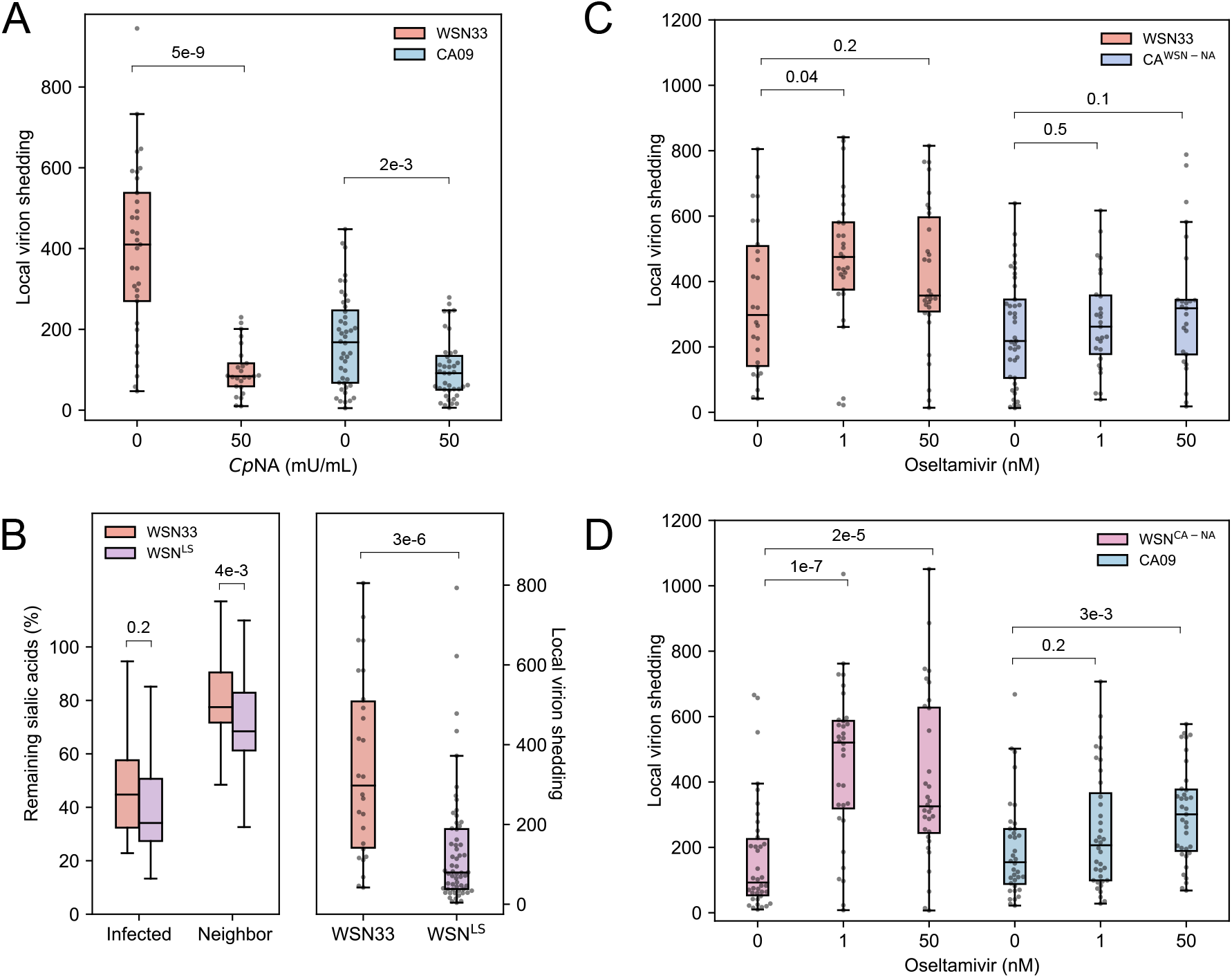
Decreasing NA activity increases local virion shedding. (A) Quantification of local virion shedding under continuous treatment with exogenous sialidase (*Cp*NA). Data is combined from three biological replicates. P-values are determined from independent t-tests. (B) *Left*: Quantification of *cis* and *trans* cleavage of Sia by WSN33 and WSN^LS^. *Right*: Quantification of local virion shedding for these strains. Data is combined from three biological replicates. P-values are determined from independent t-tests. (C) Quantification of local virion shedding for viruses with WSN33 NA (in WSN33 or CA09 backgrounds) under oseltamivir treatment. Data is combined from three biological replicates. P-values are determined from independent t-tests. (D) Quantification of local virion shedding for viruses with CA09 NA (in WSN33 or CA09 backgrounds) under oseltamivir treatment. Data is combined from three biological replicates. P-values are determined from independent t-tests.

We next perturbed the *in situ* NA activity of WSN33 by rescuing a virus (WSN^LS^) harboring a 16-residue insertion in the NA stalk that restores the length of contemporary N1 NAs and increases NA activity^45–47^. This longer-stalk NA matches CA09 NA in overall size, but preserves the catalytic domain of WSN33 NA. Consistent with prior comparisons of long- and short-stalked NAs^48–50^, our *in situ* measurements show higher *cis* and *trans* Sia depletion by this strain (Figure 4B, left). Viruses with long-stalk NA also show reduced local virion shedding relative to the parental strain (Figure 4B, right).

Although these results are consistent with a direct link between *in situ* NA activity and the extent of cellular co-infection that occurs during secondary viral spread, they do not rule out structural contributions from different NAs that may affect virus assembly. To specifically test how NA activity contributes to co-infection in the absence of genetic changes, we treated cells infected with viral strains harboring WSN33 NA and CA09 NA with varying concentrations of the NA inhibitor oseltamivir^51^. In contrast to strains harboring WSN33 NA, we found that oseltamivir treatment leads to a significant increase in local virion shedding in strains harboring CA09 NA (Figure 4C & D). This suggests that the effect of NA mutations on local virion shedding likely result from changes in enzymatic activity, as opposed to differences in virion assembly. They also highlight limitations of NA inhibitors in preventing short-range viral transmission.

### Increased cellular MOI promotes the spread of CA09 infection

High viral loads per cell may enhance infection by IAV, where the ratio of total virus particles to fully-infectious units is thought to range from ~10-100^1,52–54^. To determine how *in situ* NA activity contributes to the spread of infection, we compared the size of infection foci for cells infected with WSN33 or CA09 during multi-cycle viral spread, as well as their counterparts with swapped NAs. As an additional perturbation of NA activity, we also tested the effects of treatment with oseltamivir. For viruses harboring CA09 NA, but not those with WSN33 NA, the size of infection foci increased significantly following NA inhibition (Figure 5A & B). This follows the trends observed for these strains in local virion shedding, suggesting that increased particle counts in neighboring cells maps to increased probability of secondary infection.

**Figure 5:**
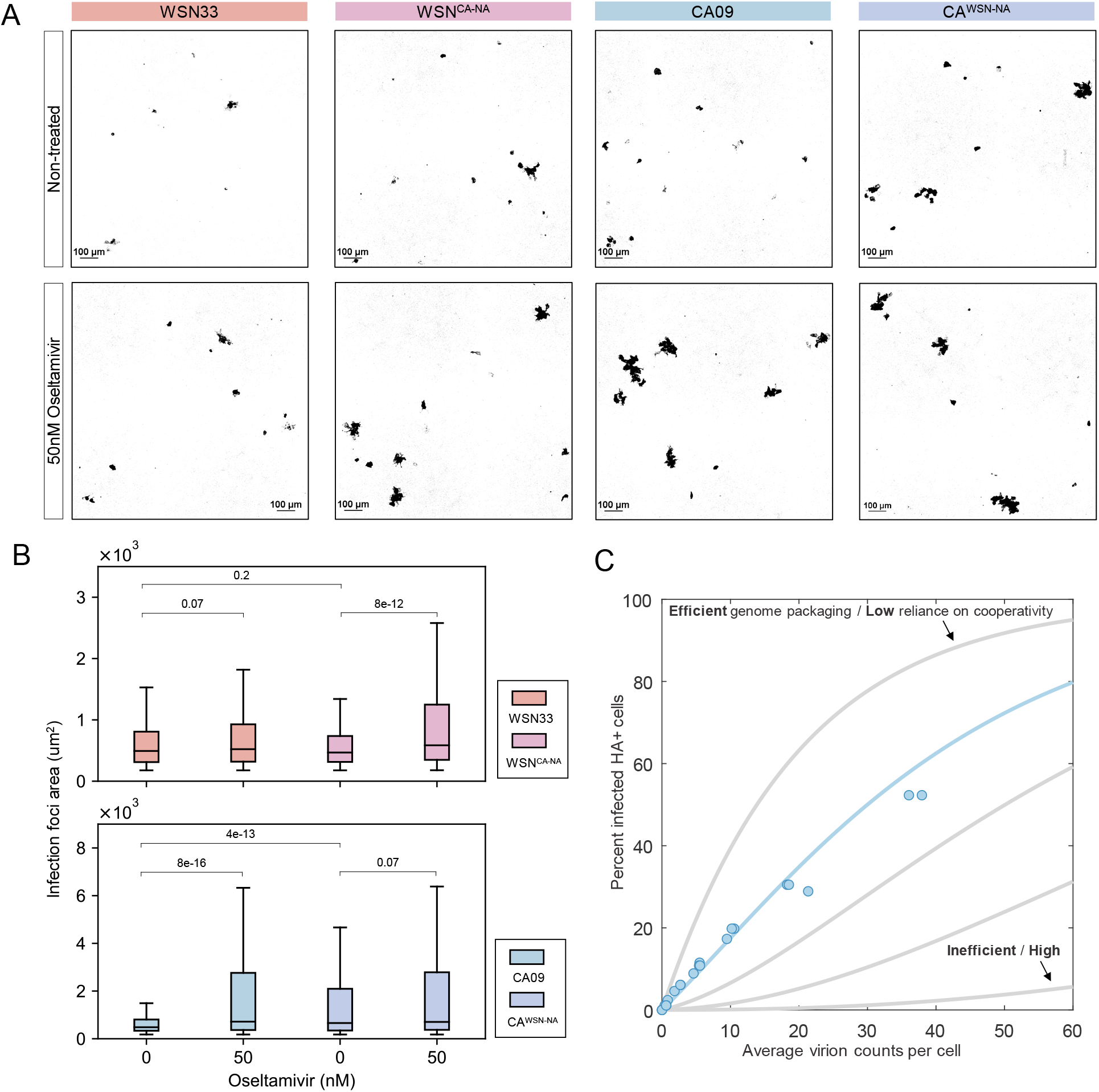
Increased cellular MOI compensates for low infectious potential. (A) Representative images of infected clusters of A549 cells at 48 h.p.i. Infected cells (shown in black) are visualized by an M2-specific Fab. (B) Quantification of the size distribution of infection foci at 48 h.p.i. Data is combined from three biological replicates. P-values are determined by Kolmogorov–Smirnov tests. (C) Relating the average virion count per cell to the probability of infection. Data for CA09 is shown as blue circles. Curves are produced from an infection model, with the best fit shown in blue. Gray curves show the effects of varying genome segment packaging probabilities.

Assuming that each infected cell is surrounded by ten nearest neighbors, our data for CA09 imply that an increase from ~10 to ~20 virions per cell (obtained by dividing the data from Figure 4D by the number of neighboring cells) significantly increases the probability of infection (Figure 5A & B). To verify if this is the case, we measured the relationship between virion counts and infection in A549 cells. We observed that the proportion of infected cells increases linearly with the average numbers of virions per cell over a wide range (Figure 5C). At an average of ~10 virions per cell, the proportion of infected cells remains modest, at around 20%. By fitting this data to an infection model (Supplementary Figure 5A, *Methods*), we find that its high linearity is consistent with a large proportion of non-infectious particles, combined with a small proportion of semi- or fully-infectious virions that deliver genome segments with relatively high efficiency (Supplementary Figure 5B). In contrast, a large proportion of semi-infectious particles with inefficient genome delivery would exhibit higher cooperativity, producing an ‘s’-shaped curve in our model (Supplementary Figure 5C). These results confirm that infection of A549 cells by CA09 is inefficient at low particle counts, but can be increased during secondary infection by attenuating NA activity.

### The predicted optimal balance between IAV surface proteins depends on infectious potential

Collectively, our results suggest a relationship between sialic acid availability, cellular MOI, and infectious potential – characteristics that will vary widely across IAV strains and their possible hosts. To broaden our investigation beyond the strains used in our experiments, we developed a simplified model that uses probabilistic virus attachment to simulate the spread of infection (Figure 6A; *Methods*). We model viral attachment using two probabilities: a *cis* binding probability describes the likelihood that the virus will attach to the initial infected cell during a single random encounter, while a *trans* binding probability describes the likelihood of virus attachment to naïve cells. These probabilities provide a framework for modeling HA-NA functional balance. NA activity tends to drive the *cis* binding probability towards zero, and may also affect the *trans* binding probability (Figures 1 & 2), while HA binding avidity will affect both (Figure 3). This model provides estimates of cellular MOI and infection across a monolayer of cells, allowing us to determine how the surface features of a viral strain (*i.e*., its *cis* and *trans* binding probabilities) are functionally related to its dependence on co-infection (the number of virions necessary for infection).

**Figure 6:**
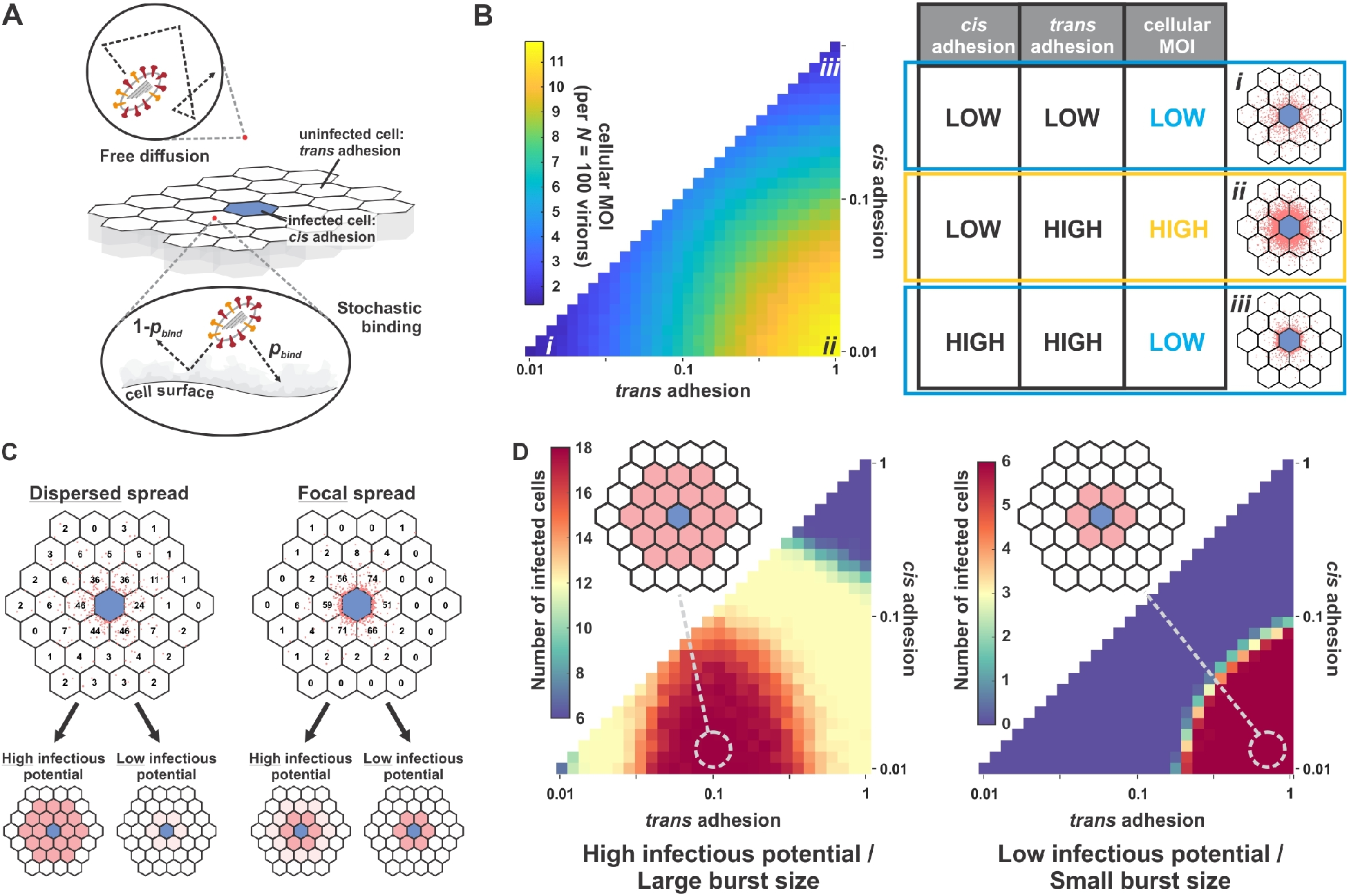
Optimal balance between IAV surface proteins depends on infectious potential. (A) Model of virus diffusion and binding to a cell monolayer. Virion attachment to the infected cell is referred to as *cis* adhesion, and attachment to uninfected cells is referred to as *trans* adhesion. During each encounter with the cell monolayer, virions bind to the cell surface or get reflected with a probability (*p*_bind_) which differs for *cis* adhesion and *trans* adhesion. (B) *Left*: Model predictions for the number of virions shed to neighboring cells (“cellular MOI”) as a function of *cis* and *trans* binding probabilities. Numerical values correspond to a scenario where the infected cell sheds a total of 100 virions. *Right*: Summary of cellular MOI as a function of *cis* and *trans* adhesion probabilities. Images correspond to conditions labeled *i* – *iii* in the plot to the left. Blue hexagons (lattice center) correspond to the infected cell; red points denote the positions of bound virions. (C) Distribution of secondary infection as a function of viral adhesion (dispersed vs. focal) and infectious potential (low vs. high). For viruses with high infectious potential, dispersed spread leads to greater secondary infection whereas focal spread produces more secondary infection when infectious potential is low. (D) Model predictions for secondary infection as a function of *cis* and *trans* adhesion. *Left*: Viruses with high infectious potential produce the greatest amount of secondary infection at intermediate *trans* adhesion (~0.1), where virion spread is more dispersed. *Right*: Viruses with low infectious potential produce the greatest amount of secondary infection at high *trans* adhesion (~1), where virion spread is focal.

Figure 6B shows the predicted number of virions per cell adjacent to the initial site of infection across a 100-fold range in *cis* and *trans* binding probabilities (0.01-1). Not surprisingly, viral load per cell is highest when viruses bind strongly to naïve cells with negligible binding to the infected cell, but can be reduced when overall adhesion is either too weak or too strong (Figure 6B, right). Predicted viral loads per cell can be used to estimate probability of infection using our measured data for CA09 (Figure 5C) or hypothetical parameters for viruses with a different dependence on co-infection. While strains requiring high viral loads for efficient infection favor focal spread in which *trans* binding probabilities are high (Figure 6C & D, right), reduced *trans* binding is more efficient when fewer virions are necessary for infection (Figure 6C & D, left). These predictions also generalize to viral burst size, with increased burst size permitting more dispersed spread for a given infectious potential, and smaller burst sizes requiring more localized spread. Overall, these results predict that optimal HA-NA balance is functionally linked to intracellular aspects of viral replication, with potential implications for viral evolution and adaptation to new hosts.

## Discussion

While the concept of HA-NA functional balance is well-established, our work provides additional insights into how the competing activities of the IAV surface proteins shape the spatial structure of viral spread, and the specific role of NA in this process. Depletion of sialic acid in both *cis* and *trans* (Figure 2) combined with genetic differences in HA binding avidity (Figure 3) collectively determine how virions spread from initial sites of infection (Figures 4–6). HA and NA activities therefore constitute a genetic mechanism through which IAV may tune the degree of cellular co-infection that occurs during multi-cycle growth.

Our work reinforces previous observations that NA cleaves Sia both in *cis* and in *trans*, and extends these observations to cell-surface NA with access to Sia on neighboring cells. The extent of cell-surface Sia depletion is similar when NA is expressed through infection or through transient transfection (Figure 2D & G bottom). This is perhaps surprising, given the potential differences in the level and duration of NA expression in these experiments. One possible explanation is that NA expressed at even modest levels is able to remove all Sia to which it has access. This may also be true for virion-associated NA, although we do not observe strong Sia depletion over the course of cellular entry (Figure 2G, top). In this case, cell-surface Sia depletion may be restricted by the rapid rate at which virions are endocytosed following attachment^55^. Further work is needed to determine how NA on the cell and viral surface separately contribute to virion release and dissemination.

The mechanistic link established here between the activities of viral surface proteins and the degree of cellular co-infection holds potential implications for virus evolution or adaptation to new hosts, where a dependence on complementation is critical for virus replication^14^. Specifically, our results suggest that strains that require higher degrees of co-infection - or that produce smaller burst sizes within a particular host - will spread most efficiently when adhesion to neighboring cells is strong, maximizing viral load per cell. While this comes at the cost of more dispersed spread, this may be beneficial to virus replication if dispersed virions are unlikely to result in productive infection. Previous work has demonstrated that the emergence of influenza viruses in new hosts is frequently accompanied by truncations in the NA stalk^56,57^. Although speculative, the link between reduced NA activity and increased cellular co-infection that we demonstrate here may provide insights into early events such as these during the adaptation of a virus to new hosts.

Additional mechanisms could further contribute to the frequency of co-infection, including transmission through tunneling nanotubes^58–60^ as well as viral aggregation^61^. Virion aggregation has been shown to promote co-infection by VSV and poliovirus^17,62^. In light of long-standing observations that IAV particles can form aggregates^63^, it is plausible that aggregation could contribute to the spread of IAV. Importantly, aggregation-dependent co-infection could operate over long distances – potentially between hosts - and could involve multiple distinct IAV genotypes, produced by different infected cells. Understanding IAV phenotypes that contribute to aggregation-dependent co-infection could improve current understanding of the constellation of viral factors that contribute to airborne transmission^64,65^.

Finally, it is important to note that understanding the cellular spread of IAV in humans remains an outstanding challenge which the design of our study does not allow us to address. *In vivo* studies in animal models demonstrate that long-range transmission of viruses within the airways are rare but potentially very important events^9,10^. Given the limited number of virions released by a single infected cell and the high proportion of these that may attach to adjacent cells, be swept away by mucociliary clearance, or neutralized by antibodies, we speculate that pioneering rounds of focal infection (such as those studied here) may be necessary before long-range dispersal becomes possible. Understanding how IAV strains differ in their dependence on focal versus dispersed spread in the human airways - and how host factors contribute to viral spread - will require additional work. The results presented here establish a framework for understanding the contributions of HA and NA to viral spread and cellular MOI in more complex environments.

## Methods

### Cells and viruses

Recombinant viruses were rescued using reverse genetics^66^. Briefly, HEK-293T and MDCK-II co-cultures were transfected with eight plasmids containing bi-directional promoters and encoding each viral genomic segment. Viral stocks were passaged and expanded using MDCK-II cells in virus growth medium comprised of Opti-MEM (Gibco), 2.5 mg/mL bovine serum albumin (Sigma-Aldrich), 1 mg/ml TPCK-treated trypsin (Thermo Scientific Pierce), and 1x antibiotic-antimycotic (Corning). The RNA of rescued virus strains was extracted with QIAmp DSP viral RNA mini kit (Qiagen), reverse transcribed and amplified with OneTaq one-step RT-PCR kit (NEB) for verification by Sanger sequencing.

Cells used in this study were purchased as authenticated cell lines (STR profiling) from ATCC and cultured under standard conditions (37 °C, 5% CO_2_) using DMEM (Gibco) supplemented with 10% fetal bovine serum (FBS) (Gibco) and 1x antibiotic-antimycotic. A549 cells for measuring virion spread and for quantifying sialic acid were maintained in cell-growth media. 36 hours prior to infection, cells were plated in an 8-chamber coverglass (Cellvis) coated with fibronectin (Sigma-Aldrich). After cells reached confluency, serum-containing media was removed and cells were washed twice with PBS (pH 7.4) (Gibco) before adding viral stocks diluted to MOI 0.003 in virus growth media. After 1 h incubation at 37 °C, cells were washed with PBS and the virus-containing media was replaced with fresh virus growth media containing fluorescent Fab fragments to monitor single or multiple round infection. For experiments in Figure 5, an additional 0.2 μg/ml TPCK-treated trypsin was included in the media to permit multi-round replication.

### Quantification of cell-surface sialic acids

Cell-surface sialic acids were labeled using aniline-catalyzed oxime ligation^40^. In comparison to labeling with *Sambucus Nigra* lectin (SNA) (Vector Laboratories) which are large and have defined preferences for specific Sia linkages, chemical labelling provides a more reproducible and quantitative measurement of cell-surface sialic acids (Supplementary Figure 3). To perform this reaction, we first prepared *solution A*: 1 mM NaIO4 (Sigma-Aldrich) dissolved in PBS supplemented with 1 mM CaCl_2_, and *solution B*: 1 mM CaCl_2_, 10mM aniline (Sigma-Aldrich), 5% FBS and 100 μM CF633 hydrazide (Sigma-Aldrich) or CF488A hydrazide (Sigma-Aldrich) in cold PBS (pH 6.5) (Teknova). Cells were cooled on ice and washed with cold PBS once before incubating with *solution A* on ice for 15 min, followed by a wash with cold PBS (pH 6.5) and incubation with *solution B* on ice for 40 min. Cells with labeled sialic acids were washed once with cold Opti-MEM and their plasma membranes were labeled with CellMask Orange (Invitrogen) at a concentration of 2.5 μg/mL in Opti-MEM for 5 min at room temperature. Cells were washed again with Opti-MEM before imaging.

### Quantification of NA activity with MUNANA

HEK-293T cells were transfected with pCAGGS plasmids containing NA sequences from WSN33, CA09, HK68, and BR07 with C-terminal Myc-tags attached via a short flexible linker (GGSEQKLISEEDL). Poly(ethylenimine) (PEI) (Polysciences) transfected cells were incubated for 48 h at 33 °C. The media was then removed, cells were washed once with PBS (pH 7.4), suspended by pipetting, and serially-diluted in PBS into a 96-well glass-bottom plate (Cellvis) as a 50 μL suspension. 50 μL of NA buffer (100 mM NaCl, 50 mM MES pH 6.5, 5 mM CaCl_2_, and 5% bovine serum albumin) supplemented with 0.25 mM 2’-(4-Methylumbelliferyl)-α-D-N-acetylneuraminic acid (MUNANA) sodium salt (Toronto Research Chemicals), was then added to each well. The plate was incubated at 37 °C for an hour before adding 100 μL stop solution (150 mM NaOH in 83% ethanol). MUNANA signal was measured with a Nikon Ti2 confocal microscope using excitation at 405 nm. In parallel with MUNANA tests, 10 μL of transfected cells from each sample were plated for imaging with the anti-myc antibody 9E10^67^ labeled with Sulfo-Cy5. Cell-surface NA expression was quantified from confocal stacks (1 μm step, collected with a 40x, 1.30-NA objective) and used to normalize activities determined from MUNANA measurements.

### Quantifying local virus shedding

Infected cells (MOI ~ 0.003) were imaged with a Nikon Ti2 confocal microscopy system using a 40x, 1.30-NA objective at 16 h.p.i. Infected cells were selected based on surface expression of HA (using 9 nM CR9114 Fab for H1N1 strains and 19 nM FI6v3 plus 19 nM H3v-47^68^ for H3N2 strains) and M2 (using a Fab fragment derived from mAb148^69^). Confocal z-stacks spanning a range of 15 μm were collected for the channel corresponding to labeled HA. Image analysis was performed on Nikon NIS Element software 5.21. Briefly, a maximum intensity projection was generated from confocal *z*-stacks, from which the body of the infected cell was identified by size and intensity and stored (*Mask A*). This mask was dilated by ~1 cell diameter to capture a concentric region surrounding the infected cell (*Mask B*). Peak detection was performed within the region between *Mask A* and *Mask B*, where bright spots corresponding to shed virions were identified and counted.

### Antibody purification and labeling

VH and VL sequences of Fabs were obtained from deposited antibody structures and cloned into backbones containing the CH1 and CL domains, respectively. Heavy chain Fab sequences were modified with a C-terminal ybbR tag for enzymatic labeling^70^ and His6-tag for affinity purification. HEK-293T cells at ~85% confluency were washed with PBS, transfected with verified clones and grown in Opti-MEM with 2% FBS for 7 days. Supernatants were collected and purified using Ni-NTA agarose (Thermo Scientific HisPur). Eluted antibodies were quantified by UV-Vis, diluted into a new buffer for enzymatic labeling (150 mM NaCl, 25 mM HEPES, 5 mM MgCl_2_) and concentrated to ~1 mg/mL by Vivaspin 20 centrifugal filter unit (MWCO 10 kDa) (Sartorius). Sfp synthase and CoA-conjugated dyes were prepared as previously described^54^ and used to perform the overnight Fab labeling reaction on ice. Excess dye was removed by PD-10 desalting columns (Cytiva). Expression and purification of full-length IgG1 antibodies (9E10 and 1G01^71^) followed a similar procedure, except using serum-free media for expression and protein A/G agarose (Thermo Scientific Pierce) for affinity purification.

### Creating polyclonal A549 SLC35A1 knock-out cells

A549 knockout cells were generated through transduction with lentivirus generated from the lentiCRISPR v2 packaging plasmid. Three sgRNA sequences were selected using CRISPR KO and the design rules described by Doench et al.^72^. These were tested in small scale via transient transfection in HEK-293Ts and the sgRNAs that yielded the highest efficiency (determined via Sia labeling) were selected for lentivirus preparation and infection into A549s. The optimal spacer sequence was 5’-<u>G</u>ACAGTGCATAAAGCAGTACA-3’ (underlined nucleotide added for efficient transcription initiation).

### Measuring virus binding avidity

To measure virion binding avidity, A549 cells with different Sia abundance were prepared by treatment with different concentrations of *Cp*NA (Roche) for 30 min at 37 °C. Simultaneously, viruses were labeled with fluorescent Fab fragments (18 nM CR9114 for H1N1, 19 nM FI6v3 plus 19 nM H3v-47 for H3N2) for 20 min at room temperature. Following *Cp*NA treatment, cells were washed with PBS twice and incubated with 100 μL virus-containing Opti-MEM at 4 °C. After incubating for 30 min, virus-containing media was removed and the cells were washed and supplemented with cold Opti-MEM. Viruses attached to the cell surface or endocytosed were imaged by the Nikon Ti2 confocal microscopy system using a 40x, 1.30-NA objective.

### Measuring infectious potential

To measure the relationship between viral particles and the probability of infection, we prepared stocks of CA09 virus in A549 cells. Viruses were concentrated approximately five-fold by centrifugation at 21,100 *g* for 30 min at 4 °C. Concentrated virus was then serially diluted and added to A549 cell monolayers. After 30 min of incubation at 37 °C, cells were washed with PBS twice and supplemented with virus growth media containing 9 nM labeled CR9114 Fab. To quantify the number of bound virions for each group, the same concentrations of viruses pre-incubated with 9 nM CR9114 Fab for 20 min were added to another group of cells and incubated for 30 min before washing off. Particle numbers were obtained by collecting confocal stacks and performing particle detection based on the maximum intensity projection. For the group incubated with untreated virus, percentage of infected cell was determined by measuring the ratio of HA-positive area to the total field of view (containing an intact cell monolayer) at ~12 h.p.i.

### Modeling infectious potential

To interpret our measurements of infection probability versus the average number of particles per cell, we developed a model that accounts for (1) virion delivery of incomplete genomes (*i.e*., semi- or fully-infectious particles); (2) virions that fail to deliver any genome segments (*i.e*., non-infectious particles); and (3) the Poissonian nature of virion attachment to cells.

#### Incomplete genomes

We define an eight-element vector, ***p***, whose elements (*p*_i_ correspond to the probabilities that a competent virion delivers segment *i* to the target cell (Supplementary Figure 5A). While each probability *p*_i_ could be distinct, for simplicity we model the probabilities as being equal across all segments. Since our data only captures cells that express HA, we assume that these cells express a minimum of 5 genomic segments: the vRNP segments (NP, PA, PB1, and PB2), along with the HA segment. This assumption is motivated by previous reports showing that secondary transcription (*i.e*., transcription from nascent vRNPs) is necessary for robust expression of other segments (Russell et al., 2018).

#### Non-infectious virions

Some fraction of virions within a population will not be capable of contributing to infection. These particles are distinct from those that enter the cells but package or deliver an incomplete genome. These particles could arise from failure to package any genomic segments (empty particles) or from failure to escape from the endosome (*e.g*., due to incomplete proteolytic activation of HA or for other reasons). To account for non-infectious virions, we define a parameter *ϕ* that describes the fraction of the viral population that contributes to infection.

#### Virion attachment to cells

Our measurements of particles per cell represent a population average which does not apply to any one particular cell within the population. To account for this, we model the distribution of particles per cell as following a Poisson distribution. If the average number of particles per cell is *N* and the fraction of these that are competent for infection is *ϕ*, then the distribution of *infection-competent virions per cell (n*) will follow:

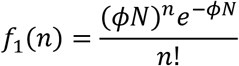

From this relationship and the segment delivery probabilities for competent virions, we can calculate infection probabilities. First, we note that the probability of failing to deliver segment *i* in *n* tries is equal to (1 – *p*_i_)^*n*^. Therefore, the probability of successfully delivering *at least one* copy of segment *i* in *n* tries is equal to 1 – (1 – *p*_i_)^*n*^. Cell-surface expression of HA indicates delivery at least one copy of at least five segments (NP, PA, PB1, PB2, and HA), the probability of which is equal to the product of the individual segment delivery probabilities:

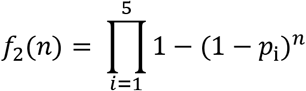

Combining the two probabilities from above, we can determine the probability of infection for a sample where the average number of particles per cell is *N*:

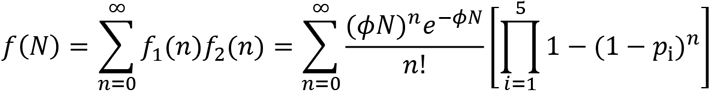

This curve can then be used to interpret our experimental measurements. In particular, this provides a means of estimating the fraction of competent virions within our sample, as well as the segment delivery probabilities (Figure 5C, Supplementary Figure 5).

### Modeling virion spread

We modeled the spread of virions from infected cells as a three-dimensional random walk above a cellular monolayer, represented by a partially-absorbing boundary. Cells within the monolayer are represented by hexagons in a lattice, whose centers are separated by 30 μm. Virions are released synchronously from a central hexagon (the infected cell) and sample a random displacement in each direction for every time step (= 1 s) of the simulation. When a virion encounters the cell monolayer, it either binds to the surface irreversibly (with a probability *p*_bind_), or reflects from the surface to continue its random walk (with a probability 1 – *p*_bind_), potentially encountering the surface repeatedly over time. We assign distinct binding probabilities to the surface of the infected cell (*cis* adhesion) and to uninfected cells (*trans* adhesion). We simulate only conditions where the *cis* adhesion probability is less than or equal to the *trans* adhesion probability, reflecting the efficient removal of Sia in *cis* during infection (Figure 2). The output of the model is a spatial distribution of bound and unbound virions that evolves over time, from which we can calculate the probability of infection using results measured for CA09 or from virions with hypothetical characteristics (Figure 6C & D).

### Statistics and replicates

Replicates referenced throughout the paper refer to biological replicates, defined as separate cultures of cells infected/ transfected/ treated individually and assayed as indicated. All statistical tests were performed in Python Scipy 1.7.3. No statistical methods were used to predetermine sample size. Statistical tests and the number of replicates used in specific cases are described in figure captions. Box plots may sometimes not show the outliers due to the limitation of y-axis.

## Acknowledgements

This work was supported by Institutional funds from Washington University in St. Louis, NIH R21 AI163985 and Burroughs Wellcome Fund CASI 1013923.

## Supplementary Figures

**Supplementary Figure 1:**
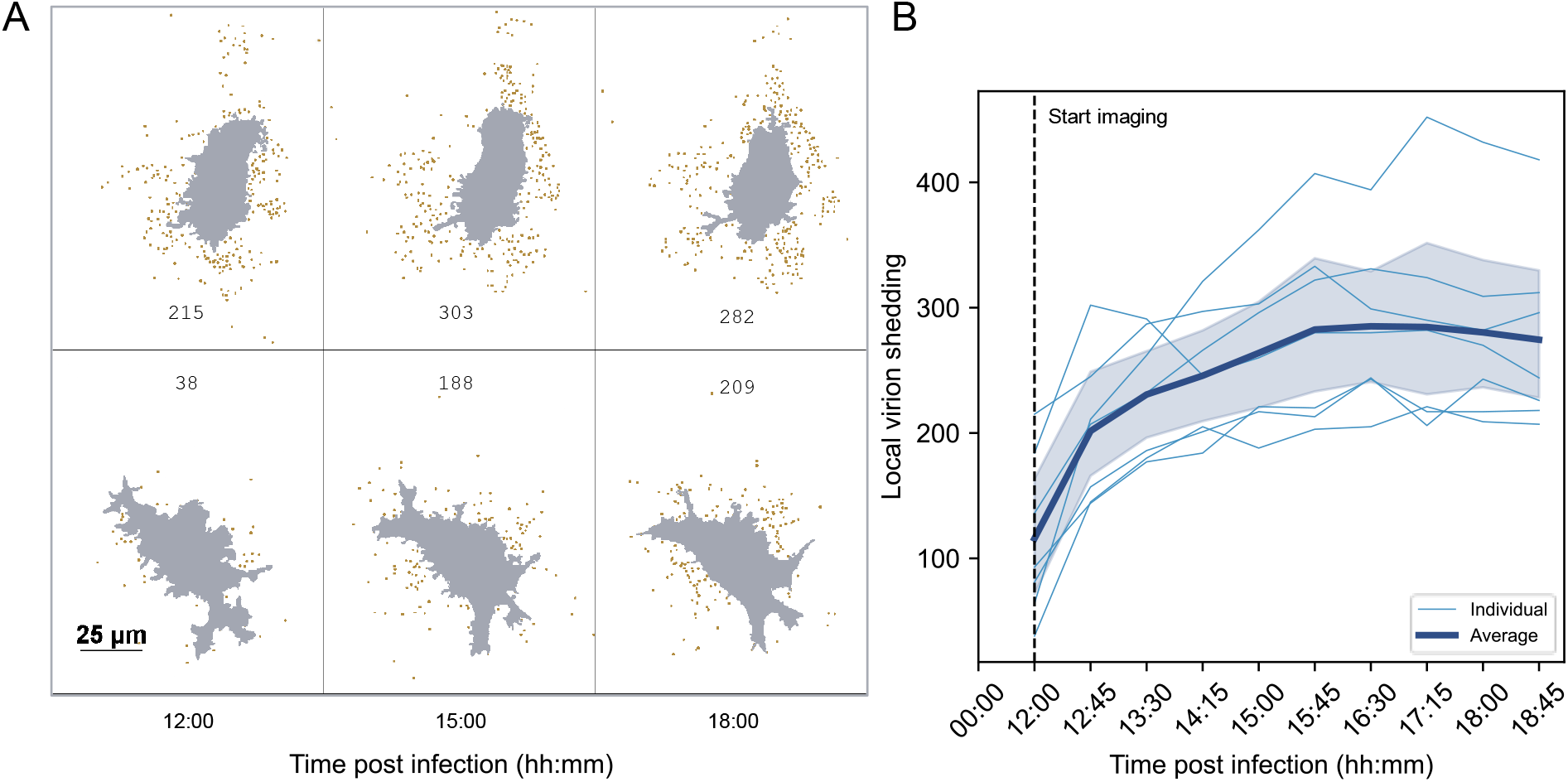
Progression of virion shedding to neighboring cells over time. (A) Monitoring local virion shedding in A549 cells infected by WSN33. Confocal stacks of infected cells are taken starting from 12 h.p.i. Gray region marks the cell body, with detected virions highlighted in gold. Cells selected for analysis show expression of both HA and M2 on the cell surface. (B) Quantification of local virion shedding compiled from time series of seven cells infected by WSN33. Shaded region represents 95% confidence interval.

**Supplementary Figure 2:**
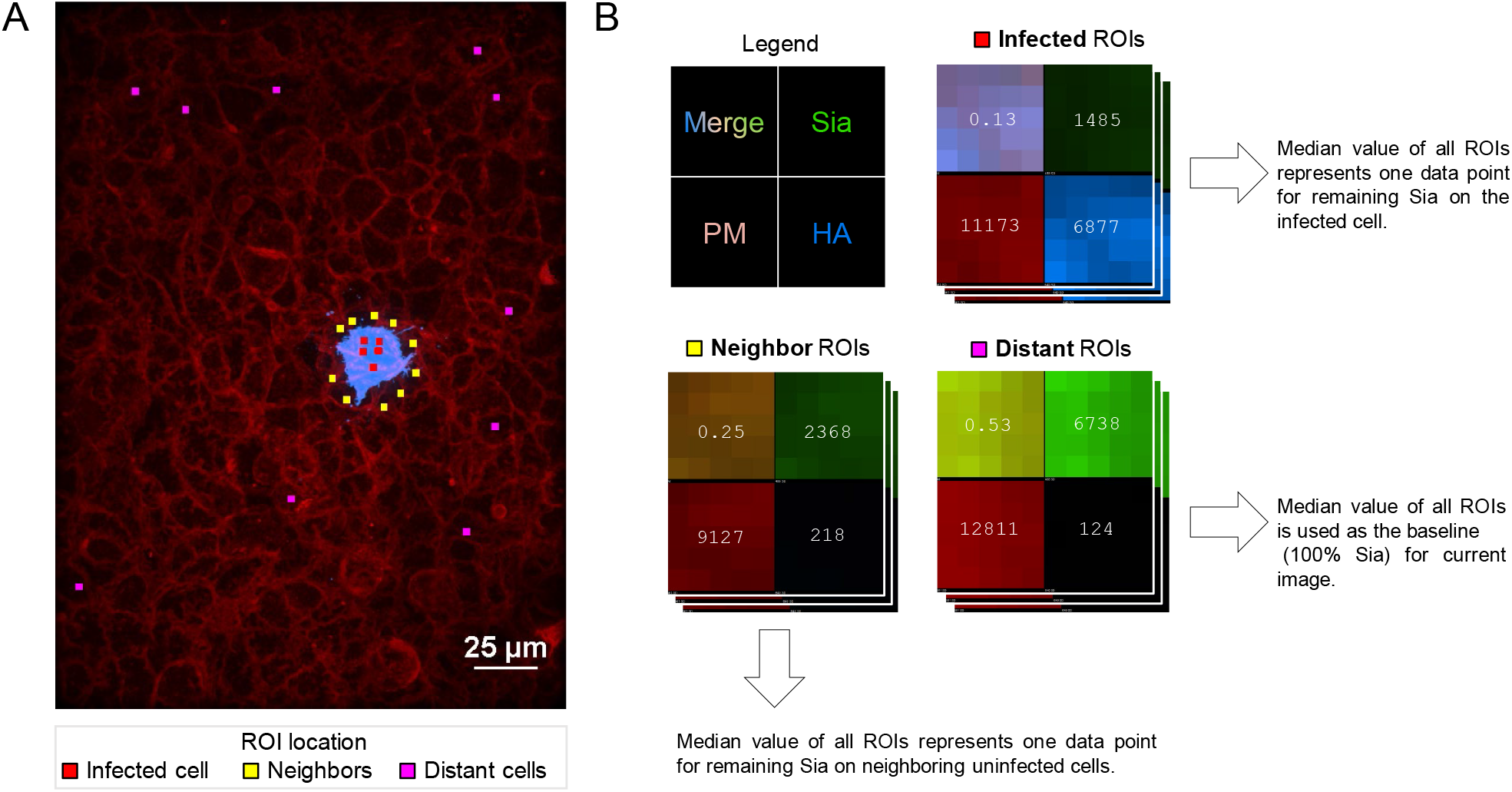
Quantifying depletion of cell-surface Sia using hydrazide coupling. (A) Image showing regions of interest (ROIs) sampled from the surface of an infected cell (HA+), uninfected neighbors, and uninfected distant cells. (B) Enlarged ROIs in split view. The number on each split channel represents the mean intensity. The number on “Merge” channel represents the Sia signal normalized by the plasma membrane (‘PM’) signal, proportional to Sia per unit membrane area. The size of each ROI is 0.55 μm by 0.55 μm.

**Supplementary Figure 3:**
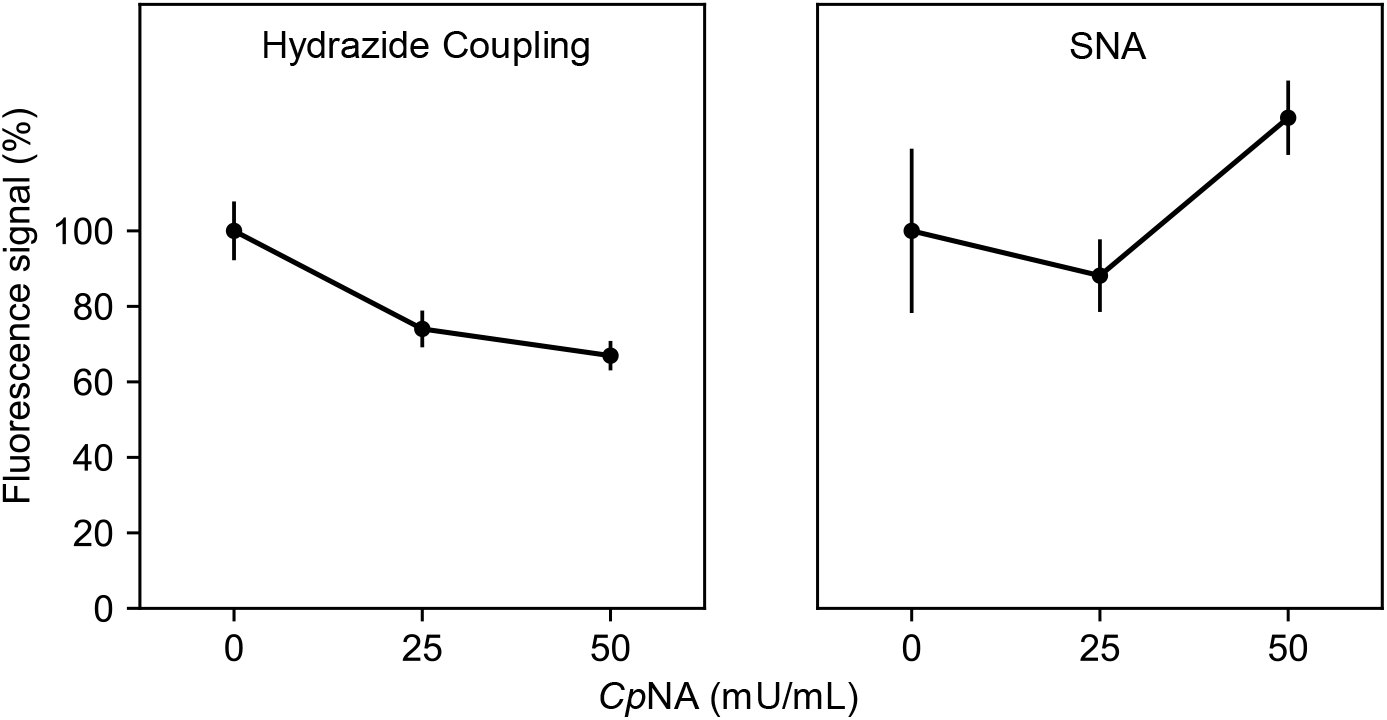
Hydrazide coupling sensitively and reproducibly reports Sia depletion by exogenous sialidase. Quantification of Sia levels on A549 cell monolayers treated with indicated concentrations of *Cp*NA for 30 min at 37°C using hydrazide coupling (left) and lectin labeling (SNA; right). Points represent mean values of three individual experiments (two for untreated group). Error bars show standard deviation.

**Supplementary Figure 4:**
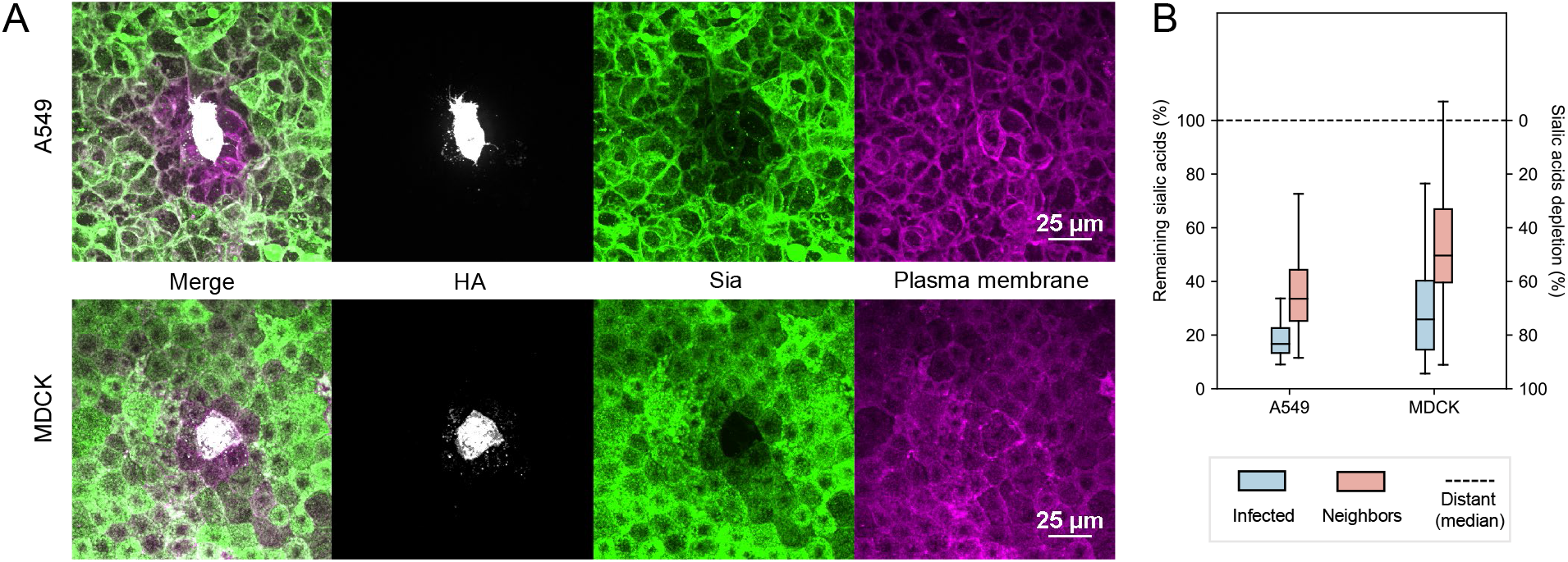
*Cis* and *trans* depletion of Sia is observed following infection of multiple cell types. (A) Images showing HA, Sia and plasma membrane in the proximity of cells infected with CA09 at MOI of 0.003. (B) Quantification of remaining Sia on infected and neighboring cells for A549 and MDCK cell lines. Data is combined from three biological replicates.

**Supplementary Figure 5:**
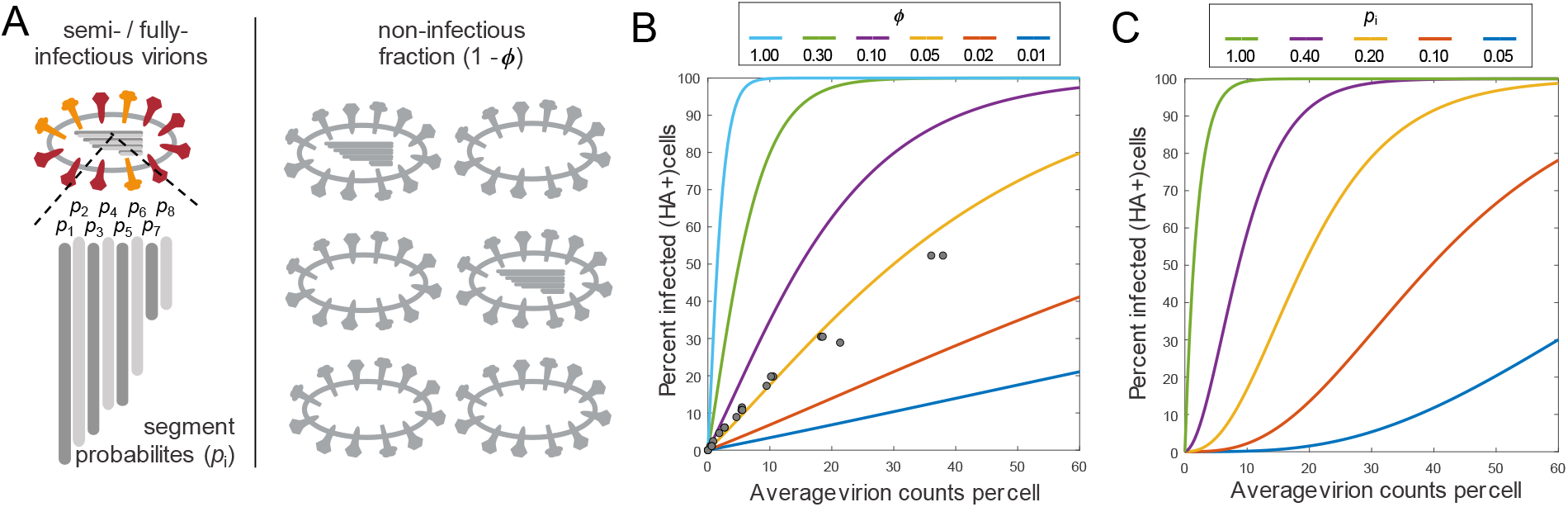
Modeling virus infection. (A) Illustration of virus populations represented by the model. Semi- or fully-infectious particles can have different probabilities of delivering each segment. The proportion of the virus population that contributes to infection is given by *ϕ*. (B) Predictions from the infection model for different segment delivery probabilities and different proportions of non-infectious virions. Results are shown for the percentage of HA+ cells. Fitting CA09 infectivity data suggests values of *p_i_* = 0.8 and *ϕ* = 0.05. (C) Measuring the percent of infected cells against average viral particles per cell under different *p*_i_ conditions where *ϕ* = 0.5.

